# Transcriptomic response of *Nitrosomonas europaea* transitioned from ammonia- to oxygen-limited steady-state growth

**DOI:** 10.1101/765727

**Authors:** Christopher J. Sedlacek, Andrew T. Giguere, Michael D. Dobie, Brett L. Mellbye, Rebecca V Ferrell, Dagmar Woebken, Luis A. Sayavedra-Soto, Peter J. Bottomley, Holger Daims, Michael Wagner, Petra Pjevac

## Abstract

Ammonia-oxidizing microorganisms perform the first step of nitrification, the oxidation of ammonia to nitrite. The bacterium *Nitrosomonas europaea* is the best characterized ammonia oxidizer to date. Exposure to hypoxic conditions has a profound effect on the physiology of *N. europaea*, e.g. by inducing nitrifier denitrification, resulting in increased nitric and nitrous oxide production. This metabolic shift is of major significance in agricultural soils, as it contributes to fertilizer loss and global climate change. Previous studies investigating the effect of oxygen limitation on *N. europaea* have focused on the transcriptional regulation of genes involved in nitrification and nitrifier denitrification. Here, we combine steady-state cultivation with whole genome transcriptomics to investigate the overall effect of oxygen limitation on *N. europaea*. Under oxygen-limited conditions, growth yield was reduced and ammonia to nitrite conversion was not stoichiometric, suggesting the production of nitrogenous gases. However, the transcription of the principal nitric oxide reductase (cNOR) did not change significantly during oxygen-limited growth, while the transcription of the nitrite reductase-encoding gene (*nirK*) was significantly lower. In contrast, both heme-copper containing cytochrome *c* oxidases encoded by *N. europaea* were upregulated during oxygen-limited growth. Particularly striking was the significant increase in transcription of the B-type heme-copper oxidase, proposed to function as a nitric oxide reductase (sNOR) in ammonia-oxidizing bacteria. In the context of previous physiological studies, as well as the evolutionary placement of *N. europaea’s* sNOR with regards to other heme-copper oxidases, these results suggest sNOR may function as a high-affinity terminal oxidase in *N. europaea* and other AOB.

**Importance:** Nitrification is a ubiquitous, microbially mediated process in the environment and an essential process in engineered systems such as wastewater and drinking water treatment plants. However, nitrification also contributes to fertilizer loss from agricultural environments increasing the eutrophication of downstream aquatic ecosystems and produces the greenhouse gas nitrous oxide. As ammonia-oxidizing bacteria are the most dominant ammonia-oxidizing microbes in fertilized agricultural soils, understanding their response to a variety of environmental conditions is essential for curbing the negative environmental effects of nitrification. Notably, oxygen limitation has been reported to significantly increase nitric oxide and nitrous oxide production during nitrification. Here we investigate the physiology of the best characterized ammonia-oxidizing bacterium, *Nitrosomonas europaea*, growing under oxygen-limited conditions.

## 1 Introduction

Nitrification is a microbially mediated, aerobic process involving the successive oxidation of ammonia (NH_3_) and nitrite (NO_2_^−^) to nitrate (NO_3_^−^) (1). In oxic environments, complete nitrification is accomplished through the complimentary metabolisms of ammonia-oxidizing bacteria (AOB) / archaea (AOA) and nitrite-oxidizing bacteria (NOB), or by comammox bacteria (2, 3). The existence of nitrite-oxidizing archaea (NOA) has been proposed, but not yet confirmed (4). Although an essential process during wastewater and drinking water treatment, nitrification is also a major cause of nitrogen (N) loss from N amended soils. Nitrifiers increase N loss through the production of NO_3_^−^, which is more susceptible to leaching from soils than ammonium (NH_4_^+^), serves as terminal electron acceptor for denitrifiers, and contributes to the eutrophication of downstream aquatic environments (5).

In addition, ammonia oxidizers produce and release nitrogenous gases such as nitric (NO) and nitrous (N_2_O) oxide during NH_3_ oxidation at a wide range of substrate and oxygen (O_2_) concentrations (6, 7). Nitrogenous gases are formed through enzymatic processes (8-13), but also by a multitude of chemical reactions that use the key metabolites of ammonia oxidizers, hydroxylamine (NH_2_OH) and NO_2_^−^ (or its acidic form HNO_2_), as the main precursors (14, 15). AOB, in particular, release NO and N_2_O either during NH_2_OH oxidation (16-21) or via nitrifier denitrification - the reduction of NO_2_^−^ to N_2_O via NO (22-25). The first pathway is the dominant process at atmospheric O_2_ levels, while the latter is more important under O_2_-limited (hypoxic) conditions (26, 27), where NO_2_^−^ and NO serve as alternative sinks for electrons generated by NH_3_ oxidation.

*Nitrosomonas europaea* strain ATCC 19718 was the first AOB to have its genome sequenced (28), and is widely used as a model organism in physiological studies of NH_3_ oxidation and NO/N_2_O production in AOB (27, 29-36). The enzymatic background of NO and N_2_O production in *N. europaea* is complex and involves multiple interconnected processes (Fig. 1). Most AOB encode a copper-containing nitrite reductase, NirK, which is necessary for efficient NH_3_ oxidation by *N. europaea* at atmospheric O_2_ levels. NirK is also involved in but not essential for NO production during nitrifier denitrification in *N. europaea* (26, 27, 29, 35), and is upregulated in response to high NO_2_^−^ concentrations (37). Moreover, two forms of membrane-bound cytochrome (cyt) *c* oxidases (cNOR and sNOR), and three cytochromes referred to as cyt P460 (CytL), cyt *c’* beta (CytS) and cyt *c*_554_ (CycA), have been implicated in N_2_O production in *N. europaea* and other AOB (12, 24, 32, 38-40). However, the involvement of cyt *c*_554_ in N_2_O production has recently been disputed (41). Finally, recent research has confirmed that the oxidation of NH_3_ to NO_2_^−^ in AOB includes the formation of NO as an obligate intermediate, produced by NH_2_OH oxidation via the hydroxylamine dehydrogenase (HAO) (20). The enzyme responsible for the oxidation of NO to NO_2_^−^ (the proposed nitric oxide oxidase) has not yet been identified (40).

**Figure 1.**
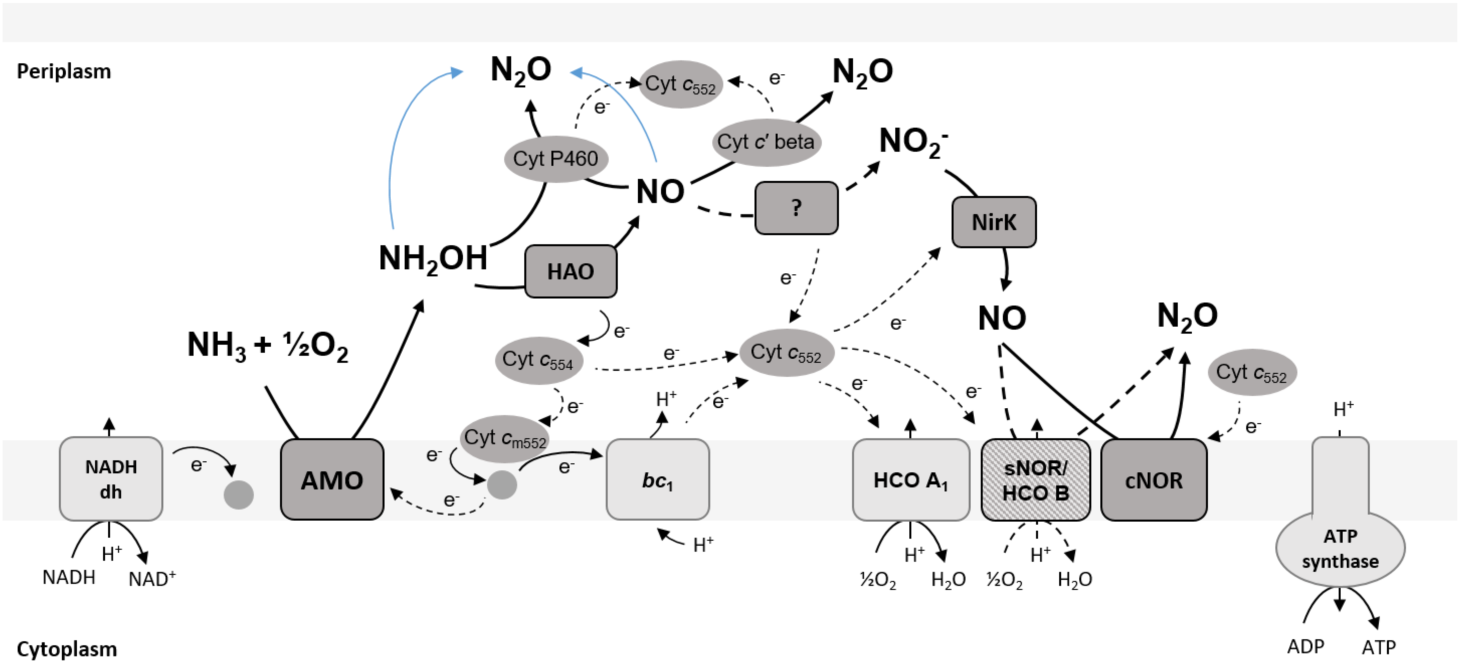
A simplified schematic of electron transport and NO/N_2_O producing pathways in *N. europaea*. Solid lines indicate confirmed and dashed lines indicate postulated reactions or electron transfer processes. Abiotic N_2_O production is indicated in blue. NADH dh - NADH dehydrogenase (complex I); AMO - ammonia monooxygenase; HAO - hydroxylamine dehydrogenase; NirK - nitrite reductase; *bc*1 - cytrochrome *bc*1 complex (complex III); HCO A1 - heme-copper containing cytochrome *c* oxidase A1-type (complex IV); sNOR/HCO B - heme-copper containing NO reductase/heme-copper containing cytochrome *c* oxidase B-type (complex IV); cNOR - heme-iron containing nitric oxide reductase.

The production of NO and N_2_O by *N. europaea*, grown under oxic as well as hypoxic conditions, has been previously demonstrated and quantified in multiple batch and chemostat culture studies (11, 12, 34, 35, 42, 43). Furthermore, recent studies have investigated the instantaneous rate of NO and N_2_O production by *N. europaea* during the transition from oxic to hypoxic or anoxic conditions (12, 35, 36). Despite this large body of literature describing the effect of oxygen (O_2_) limitation on NH_3_ oxidation and NO/N_2_O production in *N. europaea*, little attention has been paid to the regulation of other processes under these conditions. Previous studies have utilized reverse transcription quantitative polymerase chain reaction (RT-qPCR) assays to examine transcriptional patterns of specific, mainly N-cycle related genes in AOB grown under O_2_-limited conditions (34, 36, 44). To date, no study has evaluated the global transcriptomic response of *N. europaea* to O_2_-limited growth. However, research on the effect of stressors other than reduced O_2_ tension have demonstrated the suitability of transcriptomics for the analysis of physiological responses in AOB (43, 45-48).

*N. europaea* utilizes the Calvin-Benson-Bassham (CBB) cycle to fix inorganic carbon (28, 49). Whereas all genome-sequenced AOB appear to use the CBB cycle, differences exist in the number of copies of ribulose-1,5-bisphosphate carboxylase/oxygenase (RuBisCO) genes encoded, as well as the presence or absence of carbon dioxide (CO_2_) concentrating mechanisms (50-52). *N. europaea* encodes a single Form IA green-like (high affinity) RuBisCO enzyme and two carbonic anhydrases, but no carboxysome related genes (28). RuBisCO is considered to function optimally in hypoxic environments, as it also uses O_2_ as a substrate and produces the off-path intermediate 2-phosphoglycolate (53, 54). However, the effect of O_2_ limitation on the transcription of RuBisCO encoding genes and resulting growth yield in AOB is still poorly understood.

In this study, we expand upon previous work investigating the effects of O_2_ limitation on *N. europaea*, by profiling the transcriptomic response to substrate (NH_3_) versus O_2_ limitation. *N. europaea* was grown under steady-state NH_3_- or O_2_-limited conditions, which allowed for the investigation of differences in transcriptional patterns between growth conditions. We observed a downregulation of genes associated with CO_2_ fixation, as well as increased expression of two distinct heme-copper containing cytochrome *c* oxidases (HCOs) during O_2_-limited growth. Our results provide new insights into how *N. europaea* physiologically adapts to thrive in O_2_-limited environments, and identified putative key enzymes for future biochemical characterization.

## 2 Materials and Methods

### 2.1 Cultivation

*N. europaea* ATCC 19718 was cultivated at 30°C, as a batch and continuous chemostat culture as previously described (43, 48). Briefly, *N. europaea* was grown in mineral media containing 30 mmol L^−1^ (NH_4_)_2_SO_4_, 0.75 mmol L^−1^ MgSO_4_, 0.1 mmol L^−1^ CaCl_2_, and trace minerals (10 µmol L^−1^ FeCl_3_, 1.0 µmol L^−1^ CuSO_4_, 0.6 µmol L^−1^ Na_2_Mo_4_O_4_, 1.59 µmol L^−1^ MnCl_2_, 0.6 µmol L^−1^ CoCl_2_, 0.096 µmol L^−1^ ZnCl_2_). After sterilization by autoclaving, the media was buffered by the addition of 6 mL L^−1^ autoclaved phosphate-carbonate buffer solution (0.52 mmol L^−1^ NaH_2_PO_4_ x H_2_O, 3.5 mmol L^−1^ KH_2_PO_4_, 0.28 mmol L^−1^ Na_2_CO_3_, pH adjusted to 7.0 with HCl).

For steady-state growth, a flow through bioreactor (Applikon Biotechnology) with a 1 L working volume was inoculated with 2% (v/v) of an exponential phase *N. europaea* batch culture. The bioreactor was set to ‘batch’ mode until the NH_4_^+^ concentration reached <5 mmol L^−1^ (six days; Table S1). Subsequently, the bioreactor was switched to continuous flow ‘chemostat’ mode, at a dilution rate / specific growth rate (µ) of 0.01 h^−1^ (doubling time = ∼70 hours), which was controlled by a peristaltic pump (Thermo Scientific). The culture was continuously stirred at 400 rpm and the pH was automatically maintained at 7.0 ± 0.1 by addition of sterile 0.94 mol L^−1^ (10% w/v) Na_2_CO_3_ solution. Sterile filtered (0.2 µm) air, at a rate of 40 ml min^−1^, was supplied during batch and NH_3_-limited steady-state growth. Once NH_3_-limited steady-state was reached (day 7), the chemostat was continuously operated under NH_3_- limited conditions for 10 days. To transition to O_2_-limited steady-state growth, after day 16, the air input was stopped and the stirring speed was increased to 800 rpm to facilitate gas exchange between the medium and the headspace. O_2_-limited steady-state growth was achieved on day 23 as defined by the persistence of 26.4 - 31 mmol L^−1^ NH_4_^+^ and the accumulation of 22.8 – 25.5 mmol L^−1^ NO_2_^−^ in the growth medium. The culture was continuously grown under these conditions for 10 days.

Sterile samples (∼5 mL) were taken on a daily basis. Culture purity was assessed by periodically inoculating ∼100 µl of culture onto lysogeny broth (Sigma-Aldrich) agar plates, which were incubated at 30°C for at least 4 days. Any observed growth on agar plates was considered to be a contamination and those cultures were discarded. NH_4_^+^ and NO_2_^−^ concentrations were determined colorimetrically (55) and cell density was determined spectrophotometrically (Beckman) by making optical density measurements at 600 nm (OD_600_) (Table S1). Total biomass in grams dry cell weight per liter (gDCW L^−1^), substrate-consumption rate (q_NH3_), and apparent growth yield (Y) were calculated as described in Mellbye *et al.* (2016). To test for statistically significant differences in NH_4_^+^ to NO_2_^−^ conversion stoichiometry, q_NH3_, and Y between NH_3_- and O_2_-limited steady-state growth, a Welch’s t-test was performed.

### 2.2 RNA extraction and transcriptome sequencing

For RNA extraction and transcriptome sequencing, three replicate samples (40 mL) were collected on three separate days during NH_3_-limited (days 9, 10, 11) and O_2_-limited (days 28, 29, 30) steady-state growth (Fig. 2). The samples were harvested by centrifugation (12,400 × g, 30 min, 4°C), resuspended in RNeasy RLT buffer with 2-mercaptoethanol, and lysed with an ultrasonication probe (3.5 output, Pulse of 30 sec on / 30 sec off for 1 min; Heatsystems Ultrasonic Processor XL). RNA was extracted using the RNeasy minikit (Qiagen) followed by the MICROBExpress-bacteria RNA Purification Kit (Ambion/Life technologies) following the manufacturer’s instructions. Depleted RNA quality was assessed using the Bioanalyzer 6000 Nano Lab-Chip Kit (Agilent Technologies). Sequencing libraries were constructed from at least 200 ng rRNA-depleted RNA with the TruSeq targeted RNA expression Kit (Illumina), and 100 bp paired-end libraries were sequenced on a HiSeq 2000 (Illumina) at the Center for Genome Research and Biocomputing Core Laboratories (CGRB) at Oregon State University.

**Figure 2.**
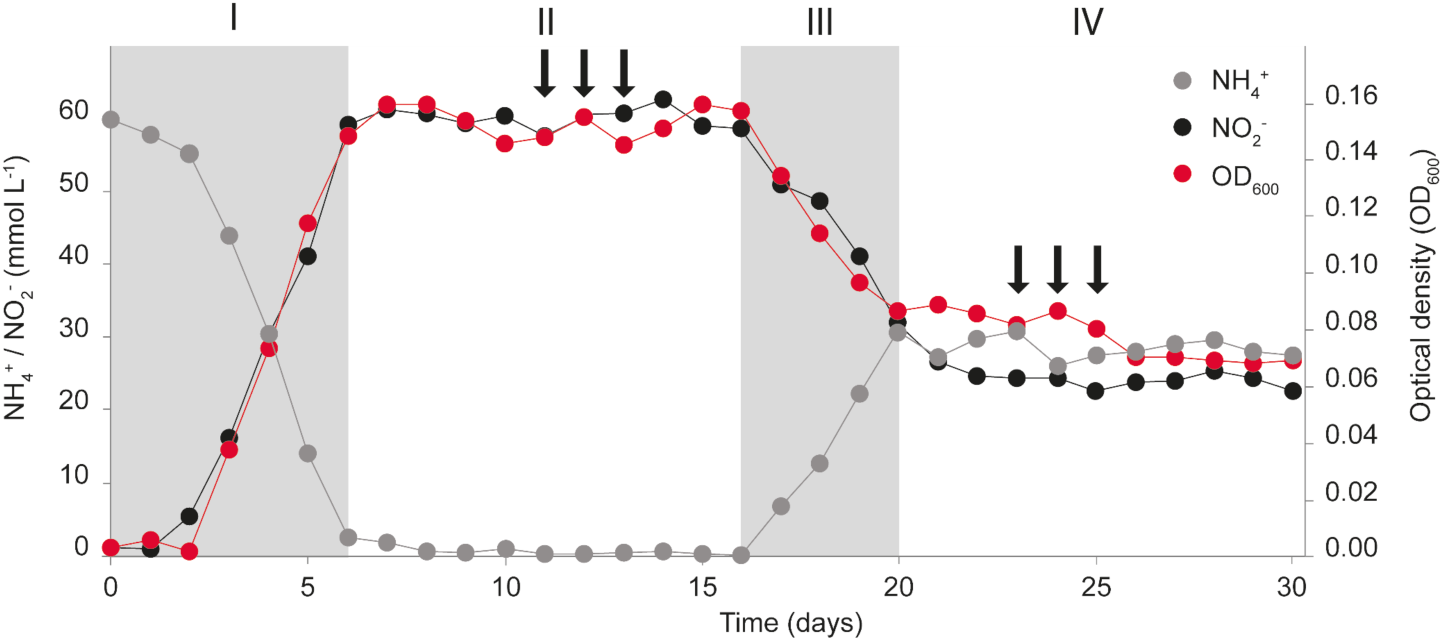
*N. europaea* culture dynamics and sampling scheme. *N. europaea* grown in a chemostat operated: in batch mode (I), under steady-state NH_3_-limited conditions as a continuous culture (II), transitioning from NH_3_-limited to O_2_-limited steady-state growth as a continuous culture (III), and under steady-state O_2_-limited conditions as a continuous culture (IV). Arrows indicate transcriptome sampling points during NH_3_-limited (days 9, 10 and 11) and O_2_-limited (days 28, 29 and 30) steady-state growth.

### 2.3 Transcriptome analysis

Paired-end transcriptome sequence reads were processed and mapped to open reading frames (ORFs) deposited at NCBI for the *N. europaea* ATCC 19718 (NC_004757.1) reference genome using the CLC Genomics Workbench (CLC bio) under default parameters as previously described (43). Residual reads mapping to the rRNA operon were excluded prior to further analysis. An additive consensus read count was manually generated for all paralogous genes. Thereafter, mapped read counts for each gene were normalized to the gene length in kilobases, and the resulting read per kilobase (RPK) values were converted to transcripts per million (TPM) (56). To test for statistically significant differences between transcriptomes obtained from NH_3_- and O_2_-limited steady-state growth, TPMs of biological triplicate samples were used to calculate p-values based on a Welch’s t-test. The more stringent Welch’s, rather than the Student’s t-Test was selected due to the limited number of biological replicates (57). Additionally, linear fold changes between average TPMs under both growth conditions for each expressed ORF were calculated. Transcripts with a p-value ≤0.05 and a transcription fold change of ≥1.5× between conditions were considered to be present at significantly different levels.

All retrieved transcriptome sequence data has been deposited in the European Nucleotide Archive (ENA) under the project accession number PRJEB31097.

## 3 Results and discussion

### 3.1 Growth characteristics

*N. europaea* was grown as a continuous steady-state culture under both NH_3_- and O_2_-limited growth conditions. During NH_3_-limited steady-state growth, the culture was kept oxic with a constant supply of filtered atmospheric air, was continuously stirred (400 rpm), and contained a standing NO_2_^−^concentration of ∼60mmol L^−1^. In contrast, during O_2_-limited steady-state growth, no additional air inflow was provided, but the stirring was increased (800 rpm) to facilitate O_2_ transfer between the headspace and growth medium. As a consequence of O_2_ limitation, the medium contained standing concentrations (∼30mmol L^−1^) of both NH_4_^+^ and NO_2_^−^ (Fig. 2, Table 1).

**Table 1:** Comparison of *N. europaea* growth characteristics and NH_4_^+^ to NO_2_^−^ conversion stoichiometry during NH_3_- and O_2_-limited steady-state growth. Letters *a* and *b* represent highly significant differences (p ≤0.01), and letters *c* and *d* represent significant differences (p ≤0.05) within parameters. Capital letters represent comparisons between 10-day periods, whereas lower case letters represent comparisons between 3-day periods.

|  | Period<br>[days] | OD<sub>600</sub><sup>1</sup> | Input NH<sub>3</sub><sup>2</sup><br>[mmol day<sup>-1</sup>] | NH<sub>3</sub> consumed<sup>1</sup><br>[mmol day<sup>-1</sup>] | Steady-state<sup>1</sup><br>NH<sub>4</sub><sup>+</sup> [mmol L<sup>-1</sup>] | Steady-state<sup>1</sup><br>NO<sub>2</sub><sup>-</sup> [mmol L<sup>-1</sup>] | N-balance<sup>1,3</sup><br>[mmol] | Ammonia oxidation rate<sup>1</sup> (q<sub>NH3</sub>)<br>[mmol gDCW<sup>-1</sup> h<sup>-1</sup>] | Apparent growth yield<sup>1</sup> (Y)<br>[gDCW mol<sup>-1</sup> NH<sub>3</sub>] |
| --- | --- | --- | --- | --- | --- | --- | --- | --- | --- |
| NH<sub>3</sub>-limited<br>growth | 7 -16 | 0.15 +0.01 | 14.4 | 14.2 +0.1 | 0.9 +0.5 | 60.1 +1.4 | 61.0 +1.7<sup>A</sup> | 24.04 +0.93<sup>C</sup> | 0.42 +0.02<sup>C</sup> |
|  | 9-11 | 0.15 ±0.004 | 14.4 | 14.2 ±0.1 | 0.9 ±0.4 | 59.1 ±1.4 | 60.0 ±1.8<sup>c</sup> | 24.73 ±0.53<sup>c</sup> | 0.40 ±0.01<sup>c</sup> |
| O<sub>2</sub>-limited<br>growth | 23-32 | 0.07 +0.01 | 14.4 | 7.5 +0.4 | 28.9 +1.5 | 24.1 +0.8 | 52.8 +1.8<sup>B</sup> | 26.44 +2.28<sup>D</sup> | 0.38 +0.03<sup>D</sup> |
|  | 28-30 | 0.07 ±0.0005 | 14.4 | 7.5 ±0.3 | 28.6 ±1.1 | 24.3 ±1.4 | 52.9 ±2.4<sup>d</sup> | 28.51 ±1.13<sup>d</sup> | 0.35 ±0.01<sup>d</sup> |

<sup>1</sup>Average values of 3 sampling days or 10 day steady-state period, ± standard deviation (Table S1).
<sup>2</sup>The NH<sub>4</sub><sup>+</sup> concentration of the influx medium (60 mmol L<sup>-1</sup>) multiplied by the influx rate (0.24 L day<sup>-1</sup>).
<sup>3</sup>Sum of effluent NH<sub>4</sub><sup>+</sup> and NO<sub>2</sub><sup>-</sup> concentrations.

During NH_3_-limited steady-state growth (days 7-16; Fig. 2), *N. europaea* stoichiometrically oxidized all supplied NH_4_^+^ to NO_2_^−^ (N-balance = 61.0 ±1.7 mmol L^−1^) and maintained an OD_600_ of 0.15 ±0.01 (Table 1). During O_2_-limited steady-state growth (days 23-32; Fig. 2), *N. europaea* was able to consume on average 31.1 ±1.5 mmol L^−1^ (51.8%) of the supplied NH_4_^+^, and maintained an OD_600_ of 0.07 ±0.01 (Table 1). A decrease in OD_600_ was expected, as the O_2_-limited culture oxidized less total substrate (NH_4_^+^), resulting in less biomass produced. The conversion of NH_4_^+^ to NO_2_^−^ was not stoichiometric during O_2_-limited growth, as only 77.5% (24.1 ±0.8 mmol L^−1^) of the NH_4_^+^ oxidized was measured as NO_2_^−^ in the effluent, resulting in an N-balance of 52.8 ±1.8 mmol L^−1^ (Table 1). The significant difference (p ≤0.01) in the N-balance between NH_4_^+^ consumed and NO_2_^−^ formed during O_2_- limited growth is in accordance with previous reports and likely due to increased N-loss in the form of NH_2_OH, NO, and N_2_O during O_2_-limited conditions (12, 35, 42, 58).

The dilution rate (0.01 h^−1^) of the chemostat was kept constant during both NH_3_- and O_2_-limited growth, and resulted in 14.4 mmol day^−1^ NH_4_^+^ delivered into the chemostat. On days (9, 10, and 11), which were sampled for NH_3_-limited growth transcriptomes, *N. europaea* consumed NH_3_ at a rate (q_NH_3) of 24.73 ±0.53 mmol gDCW^−1^ h^−1^ with an apparent growth yield (Y) of 0.40 ±0.01 gDCW mol^−1^ NH_3_. During days (28, 29, and 30) sampled for O_2_-limited growth transcriptomes, the q_NH_3 was significantly higher (28.51 ±1.13 mmol gDCW^−1^ h^−1^; p ≤0.05), while Y was significantly lower (0.35 ±0.01 gDCW mol^−1^ NH_3_; p ≤0.05). When the whole ten day NH_3_- and O_2_-limited steady-state growth periods are considered the q_NH_3 and Y trends remain statistically significant (p ≤0.05) (Table 1). Overall, NH_3_ oxidation was less efficiently coupled to biomass production under O_2_-limited growth conditions.

### 3.2 Global transcriptomic response of *N. europaea* to growth under NH_3_- versus O_2_-limited conditions

Under both NH_3_- and O_2_-limited growth conditions, transcripts mapping to 2535 out of 2572 protein coding genes (98.5%) and 3 RNA coding genes (*ffs, rnpB*, and tmRNA) were detected. Many of the 37 genes not detected here encode phage elements or transposases, some of which may have been excised from the genome in the >15 years of culturing since genome sequencing (File S1). In addition, no tRNA transcripts were detected. The high proportion of transcribed genes is in line with recent *N. europaea* transcriptomic studies, where similarly high fractions of transcribed genes were detected (43, 48). A significant difference in transcript levels between growth conditions was detected for 615 (∼24%) of transcribed genes (Fig. S1). Of these 615 genes, 435 (∼71%) were present at higher levels, while 180 (∼29%) were present at lower levels during O_2_-limited growth. Genes encoding hypothetical proteins with no further functional annotation accounted for ∼21% (130) of the differentially transcribed genes (File S1). Steady-state growth under O_2_-limited conditions mainly impacted the transcription of genes in clusters of orthologous groups (COGs) related to transcription and translation, ribosome structure and biogenesis, carbohydrate transport and metabolism, as well as energy production and conversion (Fig. 3).

**Figure 3.**
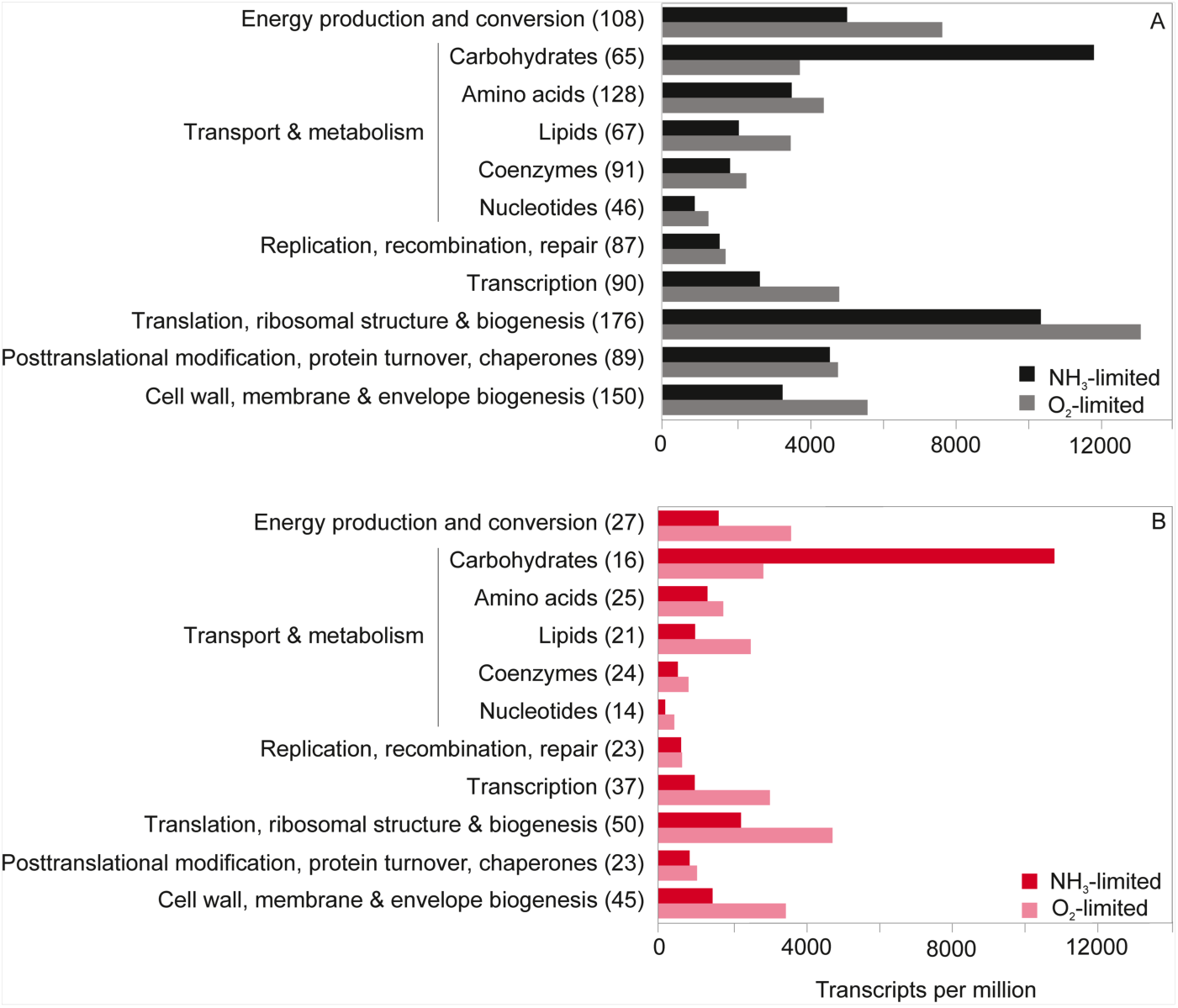
The sum of transcripts per million (TPM) for protein coding genes transcribed in given COG categories (number of transcribed genes per category is given in parenthesis) in the *N. europaea* transcriptomes: **A)** contribution and number of all transcribed genes in a given COG category; **B)** contribution and number of statistically significantly differentially transcribed genes in a given COG category.

### 3.3 Universal and reactive oxygen stress

The transcript levels of various chaperone proteins and sigma factors considered to be involved in general stress response in *N. europaea* (45) differed between NH_3_- and O_2_-limited growth with no discernible trend of regulation (Table S2, File S1). Overall, prolonged exposure to O_2_ limitation did not seem to induce a significantly increased general stress response in *N. europaea*. Key genes involved in oxidative stress defense (superoxide dismutase, catalase, peroxidases, and thioredoxins) were transcribed at lower levels during O_2_-limited growth, as expected (Table S2, File S1). Surprisingly, rubredoxin (NE1426) and a glutaredoxin family protein-encoding gene (NE2328) did not follow this trend and were transcribed at significantly higher levels (2.8- and 1.8-fold, respectively) during O_2_-limited growth (Table S2). Although their role in *N.* europaea is currently unresolved, both have been proposed to be involved in cellular oxidative stress response (60, 61), iron homeostasis (62, 63) or both.

### 3.4 Carbon fixation, carbohydrate and storage compound metabolism

There was a particularly strong effect of O_2_-limited growth on the transcription of several genes related to CO_2_ fixation (Fig. 3b). The four genes of the RuBisCO-encoding *cbb* operon (*cbbOQSL*) were among the genes displaying the largest decrease in detected transcript numbers (Fig. 4; Table S2). Correspondingly, the transcriptional repressor of the *cbb* operon (*cbbR*) was transcribed at 4.5-fold higher levels (Fig. 4, Table S2). This agrees with the previously reported decrease in transcription of the *N. europaea cbbOQSL* operon in O_2_-limited batch culture experiments (64). The reduced transcription of RuBisCO-encoding genes potentially reflects a decreased RuBisCO enzyme concentration needed to maintain an equivalent CO_2_ fixation rate during O_2_-limited growth. Since O_2_ acts as a competing substrate for the RuBisCO active site, the CO_2_ fixing carboxylase reaction proceeds more efficiently at lower O_2_ concentrations (53, 65, 66). When *N. europaea* is grown under CO_2_ limitation, the transcription of RuBisCO encoding genes increases significantly (43, 64, 67). Due to the absence of carboxysomes, *N. europaea* appears to regulate CO_2_ fixation at the level of RuBisCO enzyme concentration.

**Figure 4.**
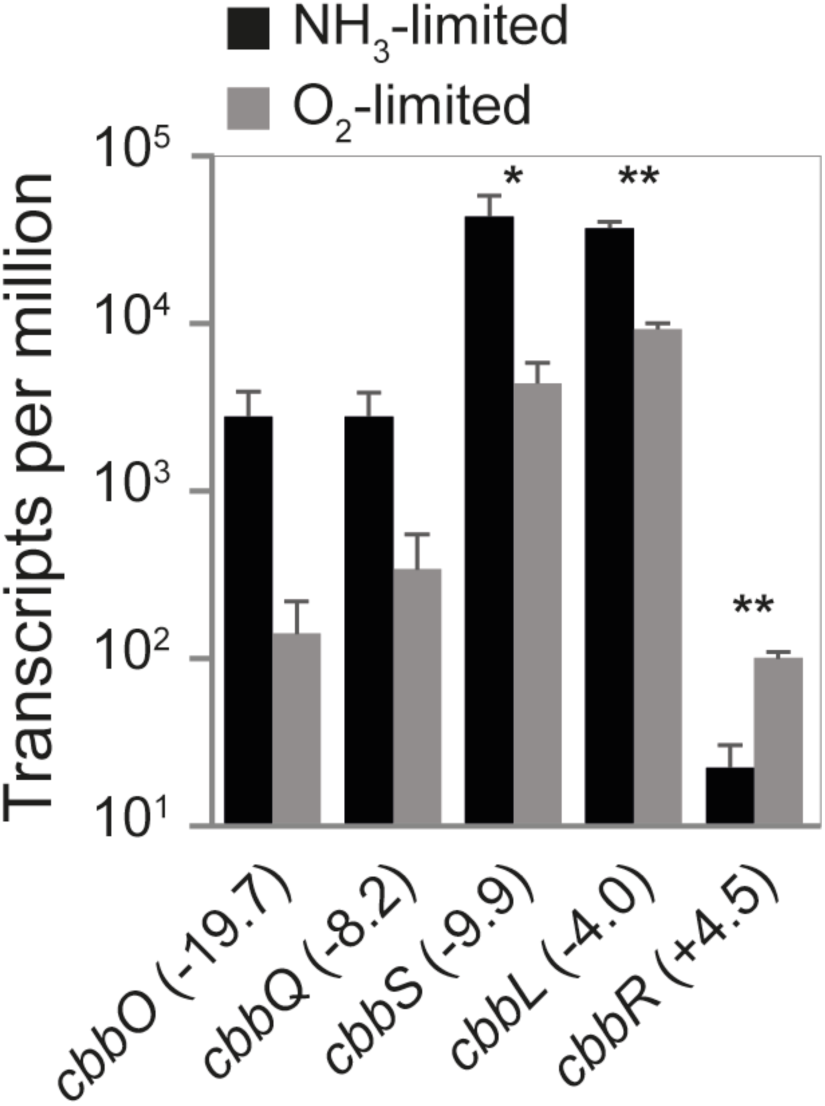
Mean TPM of all RuBisCO-encoding genes (*cbbOQSL*) and the corresponding transcriptional regulator (*cbbR*) in *N. europaea*. The fold change of gene transcription between NH_3_- versus O_2_-limited growth is given in parenthesis. Error bars represent the standard deviation between replicate samples (n=3) for each growth condition. A Welch’s t-test was used to determine significantly differentially transcribed genes (* p <0.05; ** p <0.01). For gene annotations refer to Table S2.

Genes encoding the remaining enzymes of the CBB pathway and carbonic anhydrases were not significantly differentially regulated with the exception of the transketolase-encoding *cbbT* gene (Table S2). Likewise, almost no differences in transcription were observed for the majority of genes in other central metabolic pathways (glycolysis/gluconeogenesis, TCA cycle; File S1). As the specific growth rate of *N. europaea* was kept constant during both NH_3_- and O_2_-limited growth, it is not surprising that genes associated with these core catabolic pathways were transcribed at comparable levels.

Differential transcription of polyphosphate (PP) metabolism-related genes suggests an increased accumulation of PP storage during O_2_-limited growth. Transcripts of the polyphosphate kinase (*ppk*) involved in PP synthesis were detected in significantly higher numbers (2.1-fold), while transcription of the gene encoding the PP-degrading exopolyphosphatase (*ppx*) did not change (Table S2). Indeed, *N. europaea* has previously been shown to accumulate PP when ATP generation (NH_3_ oxidation) and ATP consumption become uncoupled and surplus ATP is available (68). As the specific growth rate was kept constant throughout the experiment, PP accumulation could be a result of increased efficiency in ATP-consuming pathways, like CO_2_ fixation or oxidative stress induced repair. A decrease in the reaction flux through the energetically wasteful oxygenase reaction catalyzed by RuBisCO could result in surplus ATP being diverted to PP production.

### 3.5 Energy conservation

Genes encoding the known core enzymes of the NH_3_ oxidation pathway in *N. europaea* were all highly transcribed during both NH_3_- and O_2_-limited growth (Table S2). These included the ammonia monooxygenase (AMO; *amoCAB* operons and the singleton *amoC* gene), as well as the HAO (*haoBA*) and the accessory cyt *c*_554_ (*cycA*) and cyt c_m552_ (*cycX*) encoding genes. Due to a high level of sequence conservation among the multiple AMO and HAO operons (69), it is not possible to decipher the transcriptional responses of paralogous genes in these clusters. Therefore, we report the regulation of AMO and HAO operons as single units (Table S2). The transcript numbers of genes in the AMO operons decreased up to 3.3-fold during O_2_-limited growth, while transcripts of the singleton *amoC* were present at 1.9-fold higher levels. However, these transcriptional differences were not statistically significant. The HAO cluster genes were also not significantly differentially transcribed (Table S2).

Previous research has shown that transcription of AMO, and to a lesser extent of HAO, is induced by NH_3_ in a concentration dependent manner (70). In contrast, other studies have reported an increase in *amoA* transcription by *N. europaea* following substrate limitation (44, 71). Furthermore, *N. europaea* has been reported to increase *amoA* and *haoA* transcription during growth under low O_2_ conditions (34). However, exposure to repeated transient anoxia did not significantly change *amoA* or *haoA* mRNA levels (36). As both NH_3_- and O_2_ limitation have previously been shown to induce transcription of AMO and HAO encoding genes, the high transcription levels observed here under both NH_3_- and O_2_-limited steady-state growth conditions are not surprising.

The periplasmic red copper protein nitrosocyanin (NcyA) was among the most highly transcribed genes under both NH_3_- and O_2_-limited growth conditions (Table S2). Nitrosocyanin has been shown to be expressed at levels similar to other nitrification and electron transport proteins (72), and is among the most abundant proteins commonly found in AOB proteomes (47, 73). To date, the nitrosocyanin encoding gene *ncyA* has been identified only in AOB genomes (24), and has been proposed as a candidate for the nitric oxide oxidase (40). However, as comammox *Nitrospira* do not encode *ncyA* (2, 3, 13), and neither do all genome sequenced AOB (74), nitrosocyanin cannot be the NO oxidase in all ammonia oxidizers. In this study, a slight (1.7-fold), but not statistically significantly higher number of *ncyA* transcripts was detected during O_2_-limited growth (Table S2). This agrees with a previous study comparing *ncyA* mRNA levels in *N. europaea* continuous cultures grown under high and low O_2_ conditions (44). However, *N. europaea* performing pyruvate-dependent NO_2_^−^ reduction also significantly upregulated *ncyA*, while transcription of *amoA* and *haoA* decreased (44). Overall, there is evidence for an important role of nitrosocyanin in NH_3_ oxidation or electron transport in AOB, but further experiments are needed to elucidate its exact function.

Three additional cytochromes are considered to be involved in the ammonia-oxidizing pathway of *N. europaea*: i) cyt *c*_552_ (*cycB*), essential for electron transfer; ii) cyt P460 (*cytL*), responsible for N_2_O production from NO and hydroxylamine (39); and iii) cyt *c*’-beta (*cytS*), hypothesized to be involved in N-oxide detoxification and metabolism (24, 75). All three were among the most highly transcribed genes (top 20%) under both growth conditions (Table S2). In this study, *cytS* was transcribed at significantly lower levels (2.3-fold) during O_2_-limited growth. However, transcription of *cycB* and *cytL* were not significantly different (Table S2). While the *in vivo* function of *cytS* remains elusive, it is important to note that in contrast to *ncyA*, the *cytS* gene is present in all sequenced AOB and comammox *Nitrospira* genomes (12, 13, 52). The ubiquitous detection of *cytS* in genomes of all AOB, comammox *Nitrospira*, and in methane-oxidizing bacteria capable of NH_3_ oxidation (76), indicates that cyt *c*’-beta might play an important, yet unresolved role in bacterial aerobic NH_3_ oxidation.

### 3.6 Nitrifier denitrification

During O_2_-limited growth, *N. europaea* either performs nitrifier denitrification or experiences a greater loss of N intermediates like NH_2_OH (59) or NO (20), which leads to the observed N-imbalance between total NH_4_^+^ consumed and NO_2_^−^ produced (Fig. 2, Table 1). The Cu-containing NO_2_^−^ reductase NirK and the iron-containing membrane-bound cyt *c* dependent NO reductase (cNOR; NorBC) are considered to be the main nitrifier denitrification enzymes (24, 35). *N. europaea* NirK plays an important role in both nitrifier denitrification and NH_3_ oxidation (27), and is known to be expressed during both O_2_- replete and -limited growth (29, 30, 35). However, under O_2_-limited conditions, *nirK* was amongst the genes with the largest decrease in transcript numbers (4.2-fold) observed in this study (Fig. 5, Table S2). In *N. europaea, nirK* transcription is regulated via the nitrite-sensitive transcriptional repressor *nsrA* (30). Thus, in contrast to the *nirK* of many denitrifiers (77), *nirK* transcription in *N. europaea* is regulated in response to NO_2_^−^ concentration and not NO or O_2_ availability (31, 34, 48). The reduced O_2_ supply during O_2_-limited growth resulted in a ∼50% decrease in total NH_3_ oxidized and a ∼60% reduction in steady-state NO_2_^−^ concentration (Fig. 2, Table 1). The decrease in NO_2_^−^ concentration during O_2_-limited growth likely induced the transcription of *nsrA*, which was significantly (2.1-fold) upregulated (Fig. 5, Table S2). Therefore the large decrease in *nirK* transcription observed here may be due to the lower NO_2_^−^ concentrations and not a direct reflection of overall nitrifier denitrification activity. This hypothesis is consistent with the observation that NirK is not essential for NO_2_^−^ reduction to NO in *N. europaea*, and the presence of a not yet identified nitrite reductase in this organism. Previously, it has been shown that *N. europaea nirK* knockout mutants are still able to enzymatically produce NO and N_2_O (29, 35), even if hydrazine is oxidized by HAO instead of hydroxylamine as an electron donor (35). In addition, NO and N_2_O formation has also been observed in the AOB *Nitrosomonas communis* that does not encode *nirK* (12). The other three genes in the NirK cluster (*ncgCBA*) were differentially transcribed, with *ncgC* and *ncgB* being transcribed at lower levels (2 and 1.3-fold respectively), while *ncgA* was transcribed at a significantly higher level (2.6-fold) during O_2_-limited growth. The role of *ncgCBA* in *N. europaea* has not been fully elucidated, but all three genes have previously been implicated in the metabolism or tolerance of N-oxides and NO_2_^−^ (31).

**Figure 5.**
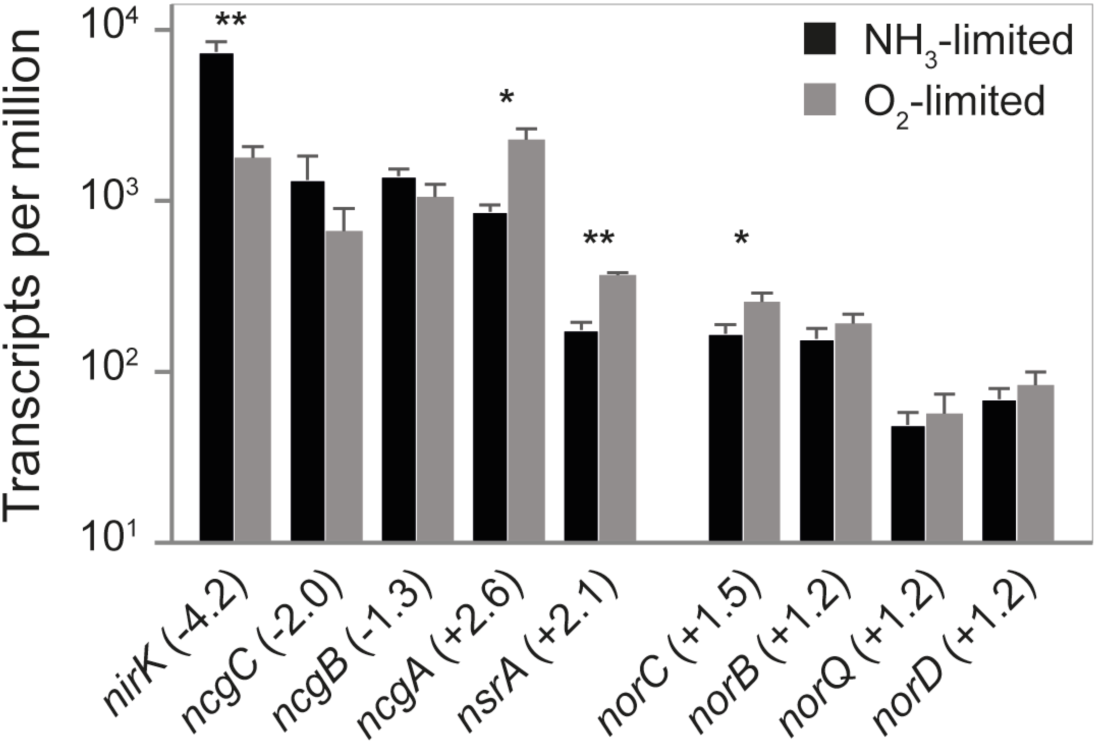
Mean TPMs of genes encoding the NirK and cNOR gene clusters in *N. europaea*. The fold change of gene transcription between NH_3_- versus O_2_-limited growth is given in parenthesis. Error bars represent the standard deviation between replicate samples (n=3) for each growth condition. A Welch’s t-test was used to determine significantly differentially transcribed genes (* p <0.05; ** p <0.01). For gene annotations refer to Table S2.

In contrast, transcripts of the *norCBQD* gene cluster, encoding for the iron-containing, cyt *c* dependent cNOR type NO reductase, were present at slightly higher (1.2- to 1.5-fold) but not significantly different levels during O_2_-limited growth (Fig. 5, Table S2). Previous research has demonstrated that in *N. europaea* cNOR functions as the main NO reductase under anoxic and hypoxic conditions (35). Interestingly, all components of the proposed alternative heme-copper containing NO reductase (sNOR), including the NO/low oxygen sensor *senC* (24) were transcribed at significantly higher levels (2.7- to 10.8-fold) during O_2_-limited growth (Fig. 6, Table S2). Therefore, it is possible that the phenotype describing cNOR as the main NO reductase in *N. europaea* (35) was a product of short incubation times and that during longer term O_2_-limited conditions sNOR contributes to NO reduction during nitrifier denitrification. Another possibility is that the increased transcription of sNOR observed here during O_2_- limited growth is primarily related to respiration and not NO reductase activity.

**Figure 6.**
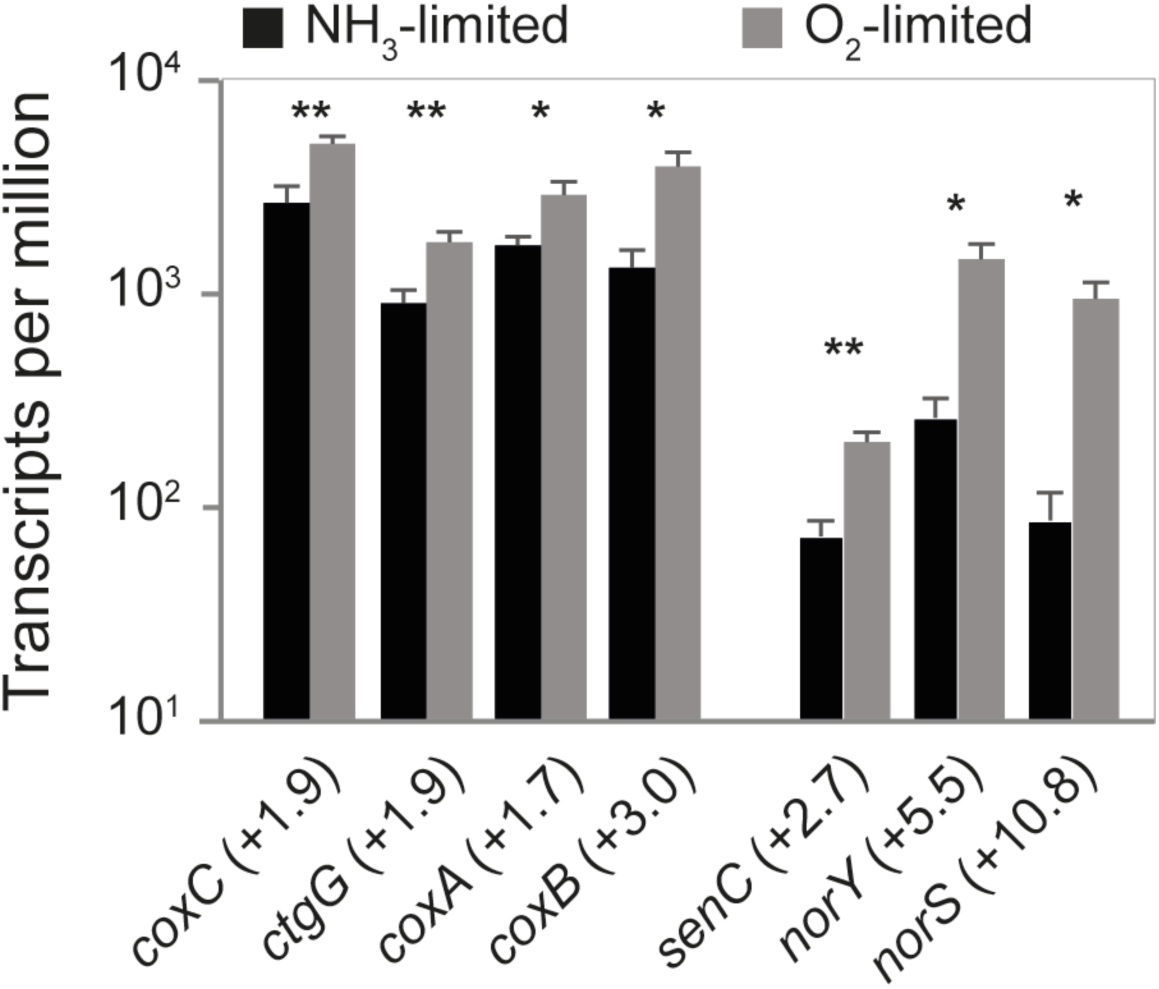
Mean TPMs of all genes encoding the A1-type and B-type HCO in *N. europaea*. The fold change of gene transcription between NH_3_- versus O_2_-limited growth is given in parenthesis. Error bars represent the standard deviation between replicate samples (n=3) for each growth condition. A Welch’s t-test was used to determine significantly differentially transcribed genes (* p <0.05; ** p <0.01). For gene annotations refer to Table S2.

### 3.7 Respiratory chain and terminal oxidases

*N. europaea* encodes a low affinity cyt *c* aa3 (A1-type) HCO, but not a high affinity *cbb*_3_-type (C-type) cyt *c* HCO encoded by other AOB such as *N. eutropha* or *Nitrosomonas* sp. GH22 (28, 50, 52). Significantly higher numbers of transcripts (1.7- to 3.0-fold) of all three subunits of the cyt *c aa*_3_ HCO and the cyt *c*-oxidase assembly gene *ctaG* were detected during O_2_-limited growth (Fig. 6, Table S2). Increased transcription of the terminal oxidase was expected, as it is a common bacterial response to O_2_ limitation (78). In addition, transcripts of all three subunits of the proton translocating cyt *bc*1 complex (Complex III) were present in higher numbers (Table S2). The NADPH dehydrogenase (Complex I) and the ATP synthase (Complex V) encoding genes were transcribed at similar levels during both growth conditions (Table S2).

As mentioned above, transcripts of both subunits of sNOR (*norSY* previously called *coxB*_*2*_*A*_*2*_), and the NO/low oxygen sensor *senC* were present at significantly higher numbers (2.7- to 10.8-fold) during O_2_-limited growth (Fig. 6, Table S2). The NO reductase function of the sNOR enzyme complex was proposed based on domain similarities between NorY and NorB (24, 32). Yet, *norY* phylogenetically affiliates with and structurally resembles B-type HCOs (79). In addition, NorY does not contain the five well-conserved and functionally important NorB glutamate residues (80), which are present in the canonical NorB of *N. europaea*. All HCOs studied thus far can reduce O_2_ to H_2_O, and couple this reaction to proton translocation, albeit B- and C-type HCOs translocate fewer protons per mol O_2_ reduced than A-type HCOs (81). Notably, NO reduction to N_2_O is a known side reaction of the A2-, B- and C-, but not A1-type HCOs (82-84). The transcriptional induction of sNOR during O_2_-limited growth reported here, as well as the high O_2_ affinity of previously studied B-type HCOs (85) indicate that sNOR might function as a high affinity terminal oxidase in *N. europaea* and possibly other sNOR-encoding AOB. Furthermore, functionally characterized B-type HCOs display a lower NO turnover rate than the more widespread high affinity C-type HCOs (82, 83). Taken together, these observations indicate that B-type HCOs, like sNOR, are ideal for scavenging O_2_ during O_2_-limited growth conditions that coincide with elevated NO concentrations, which would impart a fitness advantage for AOB growing under these conditions. Lastly, the NOR of *Roseobacter denitrificans* structurally resembles cNOR, but contains a HCO-like heme-cooper center in place of the heme-iron center of canonical cNORs. Interestingly, this cNOR readily reduces O_2_ to H_2_O, but displays very low NO reductase activity (86, 87). Therefore, in line with previous hypotheses (82, 86), the presence of a heme-copper center in NOR/HCO superfamily enzymes, such as the sNOR of *N. europaea*, may indicate O_2_ reduction as the primary enzymatic function. Notably, a recent study provided the first indirect evidence of NO reductase activity of sNOR in the marine NOB, *Nitrococcus mobilis* (88). However, further research is needed to resolve the primary function of sNOR in nitrifiying microorganisms.

## 4 Conclusions

In this study, we examined the transcriptional response of *N. europaea* to continuous growth under steady-state NH_3_- and O_2_-limited conditions. Overall, O_2_-limited growth resulted in a decreased growth yield, but did not invoke a significant stress response in *N. europaea*. On the contrary, a reduced need for oxidative stress defense was evident. Interestingly, no clear differential regulation was observed for genes classically considered to be involved in aerobic NH_3_ oxidation. In contrast, a strong decrease in transcription of RuBisCO encoding genes during O_2_-limited growth was observed, suggesting that control of CO_2_ fixation in *N. europaea* is exerted at the level of RuBisCO enzyme concentration. Futhermore, the remarkably strong increase in transcription of the genes encoding for sNOR (B-type HCO) indicates this enzyme complex might function as a high-affinity terminal oxidase in *N. europaea* and other AOB. Overall, despite lower growth yield, *N. europaea* successfully adapts to growth under hypoxic conditions by regulating core components of its carbon fixation and respiration machinery.

## Acknowledgements

We thank the Center for Genome Research and Biocomputing at Oregon State University for the sequencing services. We also thank Fillipa Sousa for helpful discussions. This work was funded by Department of Energy (DOE) award ER65192 (co-principal investigators, L.A.S-S. and P.J.B.). C.J.S., H.D., and M.W. were supported by the Comammox Research Platform of the University of Vienna. In addition, M.W. and C.J.S. were supported by the European Research Council (ERC) via the Advanced Grant project NITRICARE 294343, and C.J.S. and H.D. were supported by Austrian Science Fund (FWF) grant 30570-B29. A.T.G and D.W. were supported by the ERC Starting Grant 636928, under the European Union’s Horizon 2020 research and innovation program.

